# Neuron-type specificity of dorsal raphe projections to ventral tegmental area

**DOI:** 10.1101/2021.01.06.425641

**Authors:** Anna J. Chang, Lihua Wang, Federica Lucantonio, Maya Adams, Andrew L. Lemire, Joshua T. Dudman, Jeremiah Y. Cohen

## Abstract

The midbrain dorsal raphe (DR) and ventral tegmental area (VTA) contain two of the brains main ascending neuromodulatory transmitters: serotonin and dopamine. We studied the pathway from DR to VTA using single-cell RNA sequencing, anatomical tracing, and electrophysiology and behavior in mice. Single-cell sequencing confirmed a differential distribution of dopamine cell types between medial and lateral aspects of the VTA. This molecular diversity included differential expression of a subset of glutamatergic and serotonergic receptors. Anatomical data showed that distinct serotonergic and glutamatergic populations of DR neurons project to distinct medial-lateral locations in VTA. Physiological data showed that serotonergic neurons are positioned to excite putative dopaminergic neurons in lateral VTA on short timescales (within trial), and inhibit them on long timescales (on the next trial). Our results reveal precise anatomical specificity of DR projections to VTA, and suggest a functional role for serotonergic modulation of dopaminergic function across multiple timescales.

## Introduction

The ventral tegmental area (VTA) contains the majority of dopamine neurons that project to ventral striatum, amygdala, and frontal cortex (Morales and Margolis, 2017). Recent work has described structural and functional differences between dopamine neurons located in medial versus lateral VTA, as well as those projecting to different target sites (Gantz et al., 2018; Watabe-Uchida and Uchida, 2018). Understanding VTA dopamine heterogeneity requires knowledge of the intersection of molecular subtypes and its diverse inputs.

One of the densest afferents to dopamine neurons comes from dorsal raphe (DR) serotonin neurons, which comprise the majority of forebrain-projecting serotonin neurons. In addition, one of the densest targets of DR serotonin axons is the VTA (Azmitia and Segal, 1978; Watabe-Uchida et al., 2012; Ogawa et al., 2014; Beier et al., 2015). Serotonin concentrations are high in VTA (Saavedra et al., 1974), and serotonergic axons form synapses onto dopamine and non-dopamine neurons (Hervé et al., 1987; Van Bockstaele et al., 1994; Moukhles et al., 1997; Wang et al., 2019). Thus, the detailed study of projections from DR to VTA will be important for understanding serotonin-dopamine interactions.

As a group, DR serotonin neurons innervate almost all areas of the telencephalon and mesencephalon (Vertes, 1991; Jacobs and Azmitia, 1992). However, projection patterns from individual serotonin neurons are heterogeneous (Van Bockstaele et al., 1993; Gagnon and Parent, 2014; Ren et al., 2019). In particular, subsets of serotonin neurons project to subsets of their forebrain targets (Kiyasova et al., 2011; Commons, 2015; Kast et al., 2017; Petersen et al., 2020). These anatomical data, together with single-cell RNA sequencing experiments (Okaty et al., 2015; Huang et al., 2019; Ren et al., 2019; Okaty et al., 2020), support the hypothesis that this system may be usefully subdivided into different types (Okaty et al., 2019). In addition, electrophysiological recordings during behavioral tasks demonstrate substantial heterogeneity in the response properties of serotonin neurons and their neighbors (Nakamura et al., 2008; Ranade and Mainen, 2009; Cohen et al., 2015; Hayashi et al., 2015; Grossman et al., 2020).

The observed functional heterogeneity appears due, in part, to substantial cell-type diversity within DR. In addition to serotonin neurons (positive for tryptophan hydroxylase-2, TPH2+), DR contains non-serotonin neurons. Many of these express vesicular glutamate transporter 3 (VGLUT3+, presumably glutamatergic), and are located within approximately 200 *µ*m of the midline (Hioki et al., 2010; McDevitt et al., 2014; Qi et al., 2014; Wang et al., 2019; Cunha et al., 2020). Co-transmission is ubiquitous: a large fraction of DR VGLUT3+ neurons is also serotonergic (Shutoh et al., 2008). Dopaminergic neurons of the VTA are also characterized by diverse molecular profiles constituting several putative cell types arranged partially along a medial to lateral position (Poulin et al., 2014; La Manno et al., 2016; Poulin et al., 2020). Aside from well-characterized differences in a number of marker genes it is less clear how these differences relate to expression of serotonin and glutamate receptors (Wang et al., 2019).

Given the clear anatomical and physiological heterogeneity in DR and VTA, the pathway from DR to VTA requires more detailed analysis. A useful approach would take advantage of the known functional properties of VTA dopamine neurons. Indeed, the paranigral and parabrachial pigmented subdivisions of VTA have been well characterized anatomically and physiologically. VTA comprises multiple types of neurons, the majority of which are dopaminergic or GABAergic (Nair-Roberts et al., 2008). During many behavioral tasks, VTA dopaminergic neurons signal reward prediction errorsthe discrepancy between predicted and obtained rewards (Schultz et al., 1997; Bayer and Glimcher, 2005; Cohen et al., 2012). VTA GABAergic neurons may signal reward expectation, a quantity that their postsynaptic dopaminergic targets could use to calculate prediction errors (Cohen et al., 2012; Eshel et al., 2015).

Based on prior knowledge of the structure and function of VTA and DR, we asked three specific questions about the projections from DR to VTA: (1) Are there postsynaptic molecular differences in VTA dopamine neurons that impact DR input? (2) Is there anatomical and cell-type specificity in the DR neurons that project to VTA? (3) Do DR serotonergic projections to VTA have different effects on dopaminergic and GABAergic cell types in VTA?

To answer these molecular and anatomical questions, we used single-cell sequencing of dopaminergic neurons to identify glutamate and serotonin receptor pathway differences as a function of medial/lateral position within VTA. We then injected retrograde tracers into VTA and used immunohistochemistry to identify the molecular identity of DR neurons projecting to medial/lateral VTA targets. To test the functions of these DR→VTA projections, we selectively stimulated DR serotonergic axons in VTA while recording the activity of VTA neurons during a classical conditioning task. We used this task so that we could identify patterns of neuronal activity that have previously been mapped to neurotransmitter phenotypes (Cohen et al., 2012; Eshel et al., 2015; Tian et al., 2016; Coddington and Dudman, 2018).

We found that different neurons within DR projected to different areas within VTA. The majority of DR projections to medial VTA were TPH2+/VGLUT3+, whereas the majority of DR projections to lateral VTA were TPH2+/VGLUT3−. Serotonergic axonal stimulation had opposing effects on putative dopaminergic neurons: excitation on fast timescales (during stimulation), and inhibition on slow timescales (trials after stimulation). In contrast, we observed no clear effects of serotonergic stimulation on putative GABAergic neurons in VTA.

## Results

### Medial and lateral VTA dopaminergic neurons are transcriptionally distinct

Previous studies have shown functional differences between dopamine neurons residing in different subdivisions of VTA (Lammel et al., 2008, 2011; de Jong et al., 2019; Cai et al., 2020). Such differences can be explained by both the molecular makeup of neurons, and their presynaptic partners. Single-cell RNA sequencing experiments have revealed a diversity of gene expression products in midbrain dopamine neurons (Poulin et al., 2014; Hook et al., 2018; Saunders et al., 2018; Tiklová et al., 2019; Poulin et al., 2020). These experiments sorted neurons by dopaminergic identity using fluorescent markers, some using embryonic tissue, others using adult mice. Here, we wished to directly compare medial versus lateral VTA dopamine neurons.

We generated adult mice in which dopamine neurons were labeled with a Cre-dependent fluorescent protein (*Slc6a3* -Cre::Ai14 mice). Individual fluorescent dopamine neurons were dissociated and collected from mice expressing tdTomato in Slc6a3+ neurons in the midbrain. The medial and lateral segments of the VTA were identified (using retrograde labelling with fluorescent beads injected into nucleus accumbens core) and manually separated (schematized in Figure 1A). Collection of fluorescent cells from these tissue samples and processing for single-cell sequencing (see Methods) as in previous work (Phillips et al., 2019) yielded a sample of medial (*n* = 359) and lateral (*n* = 363) dopamine neuron transcriptomes. We first took normalized counts for all transcripts and projected each cell into two dimensions using t-distributed stochastic neighbor embedding (t-SNE; similar results with UMAP algorithm) to examine differences across samples (Figure 1A–C). As expected, we found modest gross differences between the subregions indicating similar overall expression patterns.

**Figure 1.**
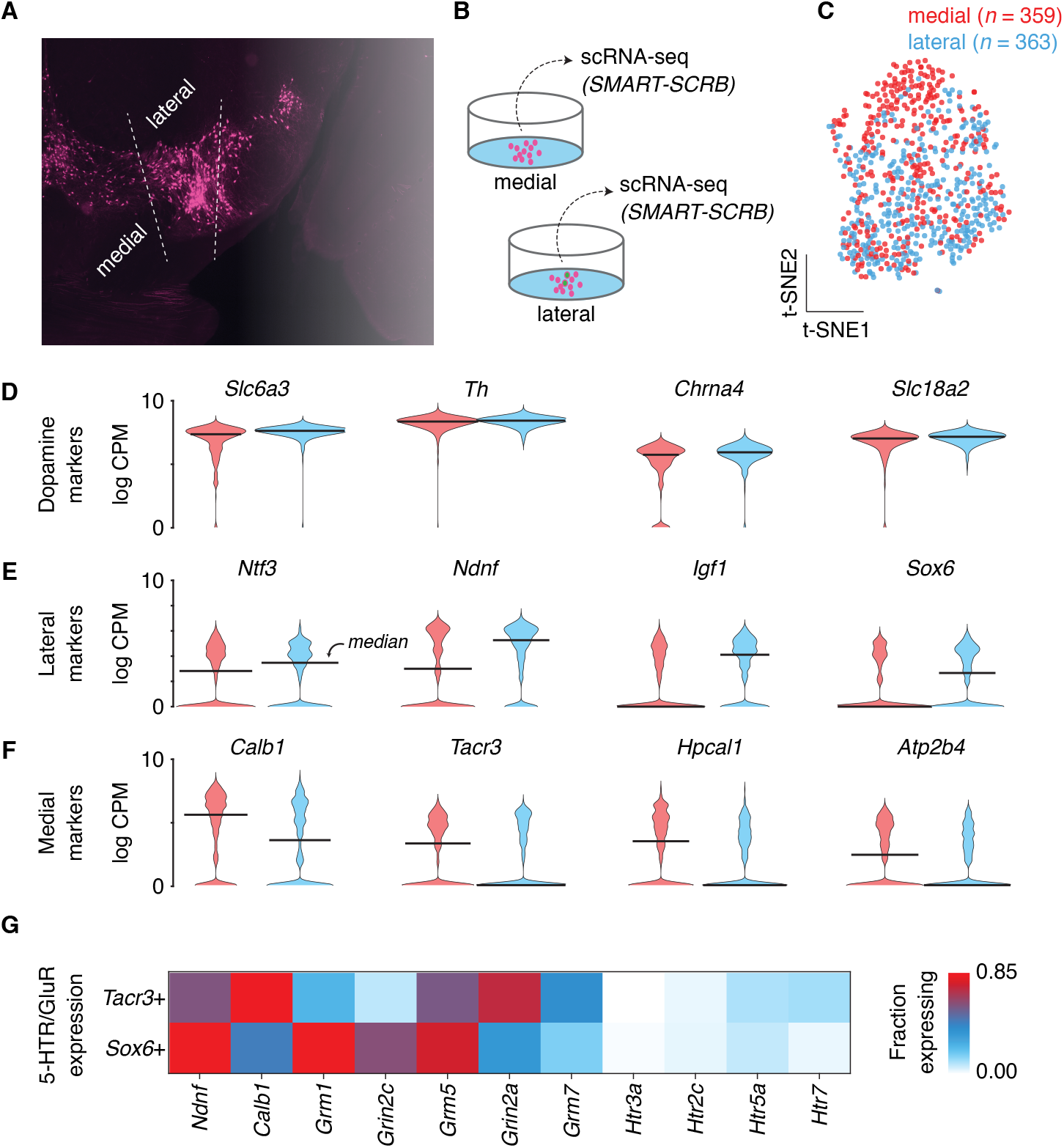
Single-cell RNA sequencing of medial versus lateral VTA dopamine neurons. (A) Coronal-plane image of VTA, showing Slc6a3+ neurons (dopamine neurons), with schematic positions of cuts to separate medial from lateral dopamine neurons. (B) Schematic of manual selection and RNA sequencing of medial versus lateral dopamine neurons. (C) t-SNE plot showing higher density of medial versus lateral dopamine neuron transcripts in the upper part of the cluster. (D-F) log CPM in medial versus lateral dopamine neurons, for four classes of transcripts. (G) Fraction of Tacr3+ (medial) and Sox6+ (lateral) dopamine neurons expressing 11 transcripts.

We next evaluated the expression of canonical dopamine neuron markers including the dopamine transporter (*Slc6a3*), tyrosine hydroxylase (*Th*), alpha-4 subunit of acetylcholine receptors (*Chrna4*), and vesicular monoamine transporter (*Slc18a2*; Figure 1D). In all cases we found robust transcripts detected and little difference across medial and lateral VTA populations, confirming similar cell health and sample preparation selecting for single dopaminergic neurons. Recent work has identified a number of genes enriched in medial and lateral VTA populations (Poulin et al., 2020). We next asked whether our medial and lateral populations were indeed enriched for marker genes previously suggested by qPCR (Poulin et al., 2014) or with improved spatial resolution via *in situ* hybridization (La Manno et al., 2016). We chose a set of 4 genes enriched in subpopulations of dopamine neurons found preferentially in lateral (*Ntf3, Ndnf, Igf1, Sox6*; Figure 1E) or medial (*Calb1, Tacr3, Hpcal1, Atp2b4*; Figure 1F) VTA. In all cases we found that our single cell sequencing recapitulated the expected lateral or medial enrichment of molecular subtypes expected.

Using these medial/lateral markers we next focused on identifying differential expression of glutamate and serotonin receptor genes presumably related to the major functional classes of DR neurons. To select for the most representative medial and lateral genes we considered the subset of single cells expressing either *Tacr3* or *Sox6*, but not both. (As in previous datasets we found “intermediate” cells expressing combinations of markers; Phillips et al., 2019.) We found several genes that were expressed above threshold in a differential fraction of individual Sox6+ or Tacr3+ neurons (Figure 1G). For example, a much larger fraction of Sox6+ (lateral) dopamine neurons expressed high levels of the metabotropic glutamate receptor *Grm1*. In contrast, the NMDA receptor subunit, *Grin2a*, was expressed in a larger fraction of Tacr3+ (medial) dopamine neurons. We observed low levels of expression of serotonin receptors in general, consistent with previous whole brain sequencing datasets (Saunders et al., 2018), although there were modest differences including a larger (albeit small) fraction of Sox6+ (lateral) VTA dopamine neurons expressing *Htr3a* (Wang et al., 2019) and a larger fraction of Tacr3+ (medial) VTA dopamine neurons expressing *Htr7*. For comparison we also provide a comprehensive mapping of all metabotropic receptors across medial and lateral VTA samples (Figure 1–figure supplement 1).

Thus, single-cell sequencing of VTA dopamine neurons segregated into medial and lateral aspects recapitulated well known molecular markers. Analysis of subsets of dopamine neurons characterized by exclusive expression of medial/lateral molecular markers revealed differential expression of a subset of glutamate and serotonin receptors. Given the heterogeneity of individual DR neurons, which can either be serotonergic or co-release glutamate, that target VTA densely we turned to retrograde tracing combined with immunohistochemistry to dissect the medial-VTA-versus lateral-VTA-projecting DR cells.

### Different DR serotonin neurons project to medial versus lateral VTA

The cell types in DR that project to VTA have been studied using retrograde tracing combined with immunohistochemistry or *in situ* hybridization. However, previous studies used relatively large retrograde tracer injections into VTA, potentially pooling subpopulations of neurons. We injected fluorescent microspheres (“retrobeads”) in VTA of 8 mice, varying the location of injection in the medial-lateral axis (Figure 2A). These retrobeads were taken up by axonal afferents in VTA and transported retrogradely to their somata in DR. Across mice, we identified DR neurons containing retrobeads. Retrogradely-labeled neurons were found in multiple subdivisions: dorsal core, ventral core, and lateral wings. Most neurons found in the lateral wings were located ipsilateral to the VTA injection site. Because the shape of DR changes in the anterior-posterior axis, we localized retrogradely-labeled neurons in four coronal sections in each mouse. Relative to Bregma, these were at −4.5 mm, −4.6 mm, −4.7 mm, and −4.8 mm. Across the 8 experiments, the labeled DR neurons tended to be farther from the midline and more dorsal (closer to the aqueduct) when the VTA injection site was more medial (Figure 2B). These correlations were larger in more posterior sections of DR (Figure 2C).

**Figure 2.**
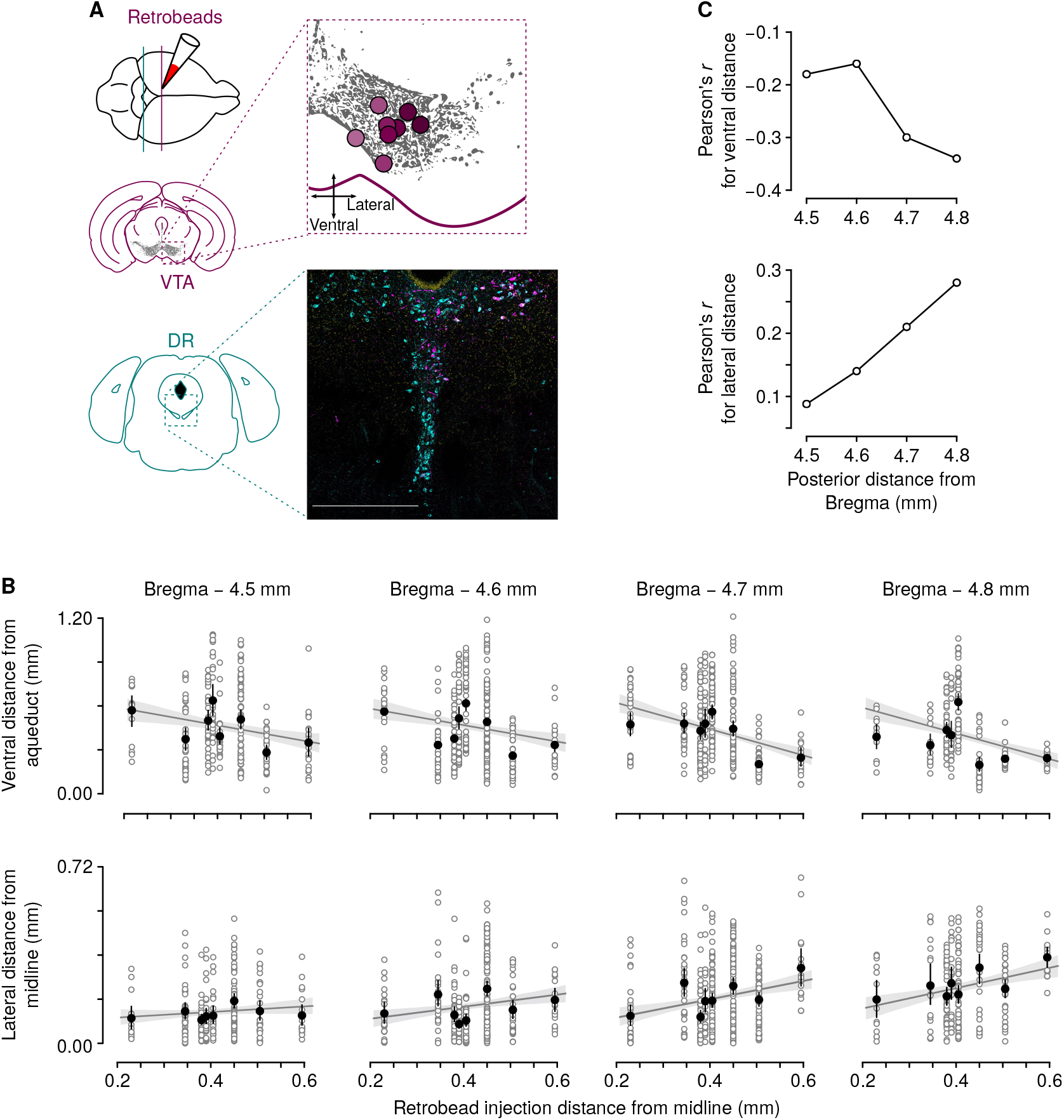
Locations in DR of retrogradely-labeled cells from VTA. (A) Schematic of retrobead injections at different mediallateral axis sites in VTA. Points correspond to estimates of the ventral-most spread of retrobeads. Scale bar: 100 *µ*m. (B) Relationships between ventral distance from aqueduct (top row) and lateral distance from midline (bottom row) as a function of estimated distance of retrobead injection from midline, at four coronal planes relative to Bregma. (C) Correlation coefficients aggregated from the plots in (B).

We found distinct topography of DR→VTA projections, roughly organized in a medial-to-medial and lateral-to-lateral fashion. Given this structure, are there also different cell types that project from DR to VTA? That is, do the DR neurons projecting to lateral VTA comprise different phenotypes than those projecting to more medial VTA? We combined the retrograde tracing described above with immunohistochemistry to detect the two predominant cell types in DR: serotonergic (i.e., TPH2+) and glutamatergic (i.e., VGLUT3+; Figure 3).

**Figure 3.**
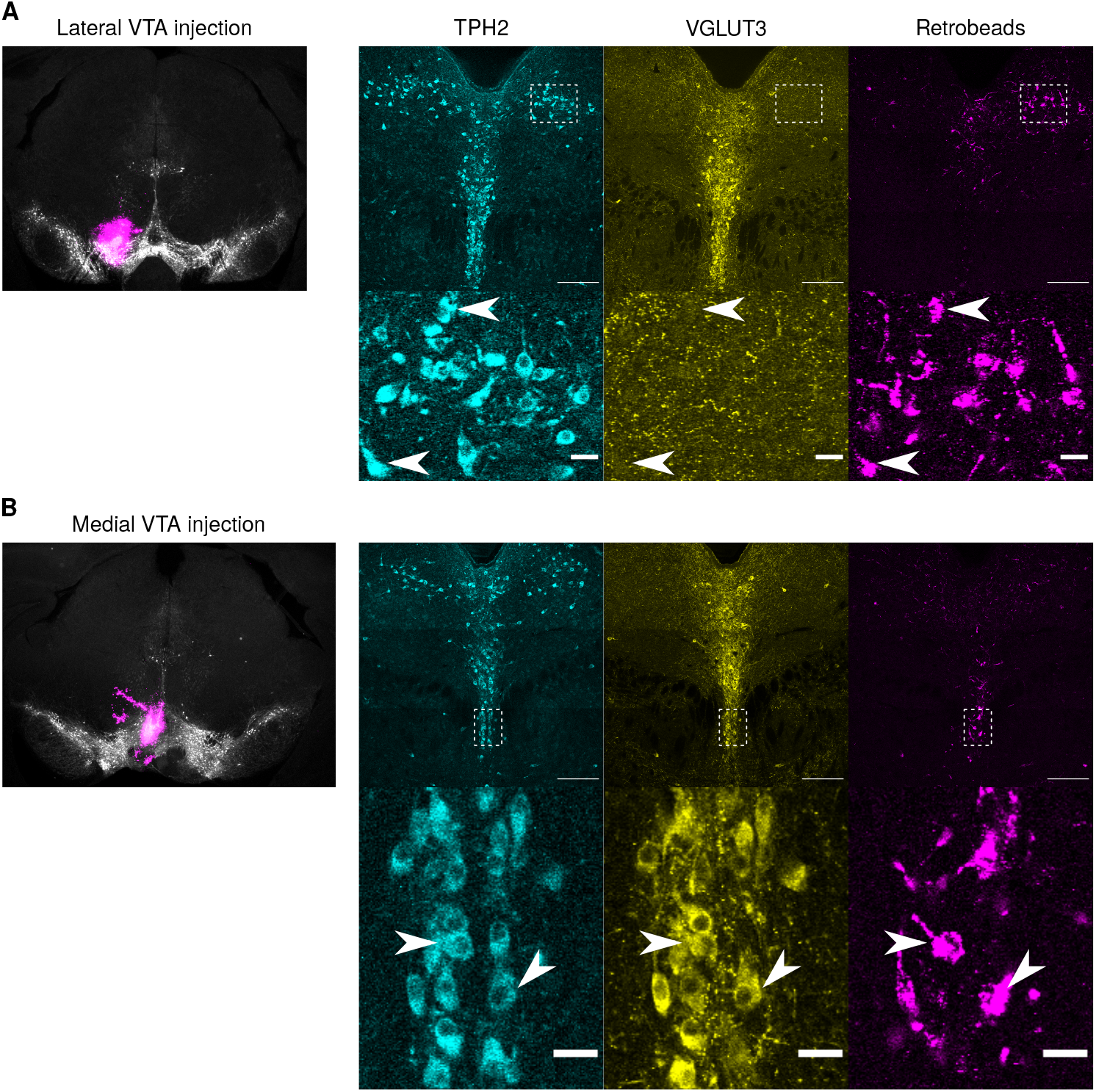
Representative medial and lateral injections yield distinct retrogradely-labeled DR neurons. (A) Left: coronal section of TH immunostaining from a mouse with a relatively lateral injection of retrobeads in VTA. Right: TPH2 and VGLUT3 immunostaining, and retrobeads, in lower magnification (top) and higher magnification (bottom, from the dashed box in the top row). Arrowheads indicate examples of retrobead-containing neurons that were TPH2+ and VGLUT3−. Scale bars: *µ*m (upper) and *µ*m (lower). (B) The same data as in (A) but from a mouse with a relatively medial injection of retrobeads in VTA. Arrowheads indicate examples of retrobead-containing neurons that were TPH2+ and VGLUT3+.

We defined 4 phenotypes of DR neurons according to the presence or absence of these two markers: TPH2+/VGLUT3−, TPH2−/VGLUT3+, TPH2+/VGLUT3+, or TPH2−/VGLUT3−. In agreement with previous work, we found that TPH2+ neurons were distributed in all DR subdivisions, while VGLUT3+ neurons were mostly located closer to the midline (Fu et al., 2010; Ren et al., 2018; Huang et al., 2019; Okaty et al., 2020). We found that VTA projections from the 4 DR phenotypes varied along the medial-lateral axis in VTA (Figure 4A). In particular, DR inputs to medial VTA were largely TPH2+/VGLUT3+ neurons, while DR inputs to lateral VTA were largely TPH2+/VGLUT3− neurons (Figure 4).

**Figure 4.**
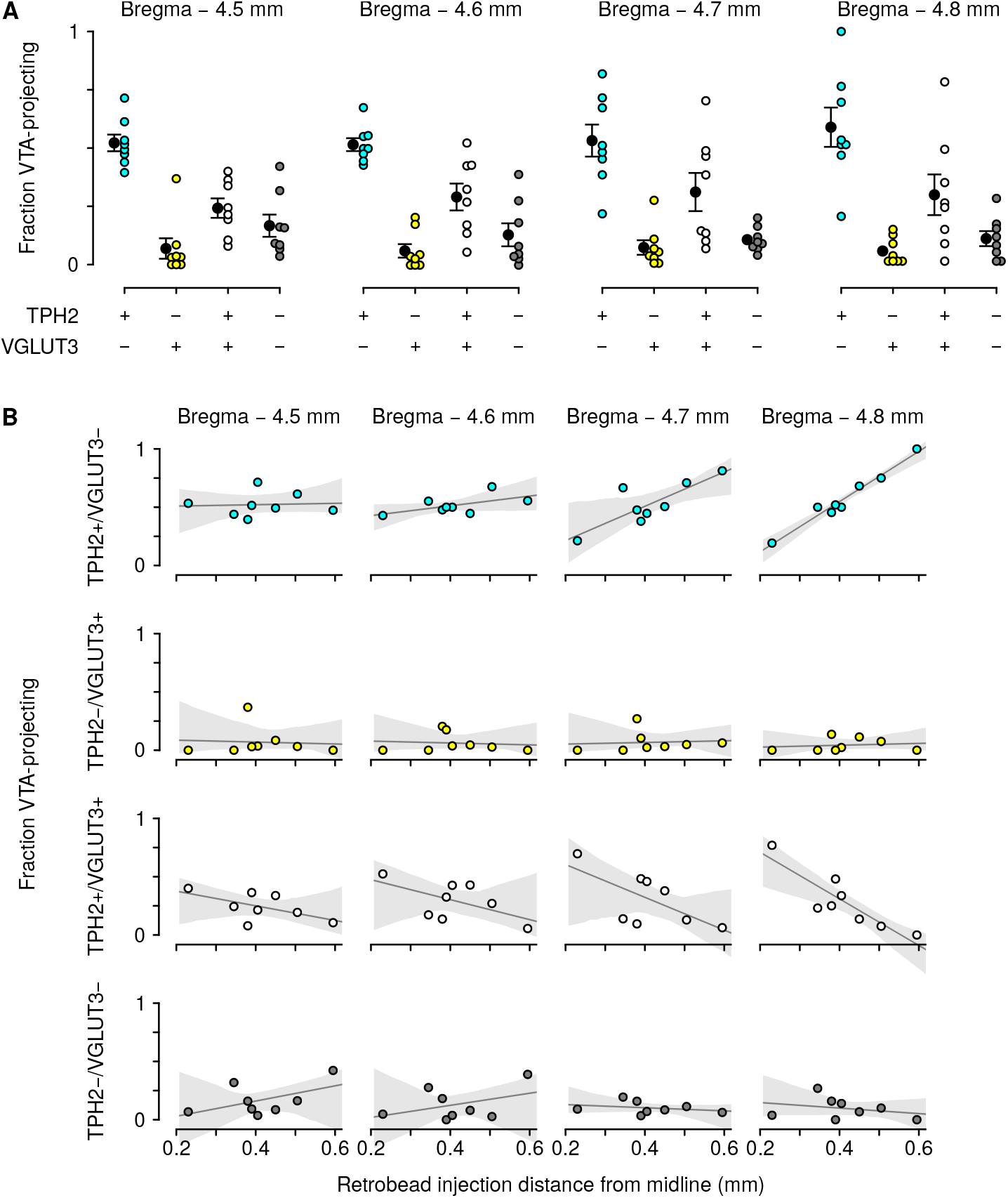
Population data of DR phenotypes that were retrogradely-labeled from VTA. (A) Fraction of VTA-projecting DR neurons at four coronal planes, across all 8 mice, that were of each of the 4 phenotypes: TPH2+/VGLUT3, TPH2/VGLUT3+, TPH2+/VGLUT3+, or TPH2/VGLUT3. (B) Relationships between fraction of VTA-projecting DR neurons of each phenotype as a function of retrobead injection distance from midline.

### Serotonergic stimulation mimics RPEs by driving fast excitation and slow inhibition in putative dopamine neurons

What are the possible functions of serotonergic inputs to VTA neurons? There are several proposals in the literature. Recent studies have focused on short-timescale effects, measuring VTA neuron activity or dopamine release in nucleus accumbens during DR neuron stimulation (Gervais and Rouillard, 2000; Qi et al., 2014; Wang et al., 2019). Theoretical studies have proposed that serotonin could act as an “opponent” to dopamine (Daw et al., 2002; Boureau and Dayan, 2011), for example, by subtracting away activity from dopaminergic neuron evoked responses (Daw et al., 2002). This could provide a mechanism for serotonergic modulation of learning in dopamine neurons.

We wished to measure activity of VTA neurons while manipulating their DR afferents in awake, behaving animals. Thus, we trained thirsty, head-restrained mice on a classical conditioning task, in which three different olfactory stimuli (conditioned stimuli, CS) predicted three outcomes: no reward, a small reward (1.5 *µ*l of water), or a large reward (4 *µ*l of water; Figure 5A). Each CS was presented for 1 s, followed by a 1-s delay and then the outcome. We used this task to replicate the clustering of VTA neurons previously observed (Cohen et al., 2012; Tian et al., 2016).

**Figure 5.**
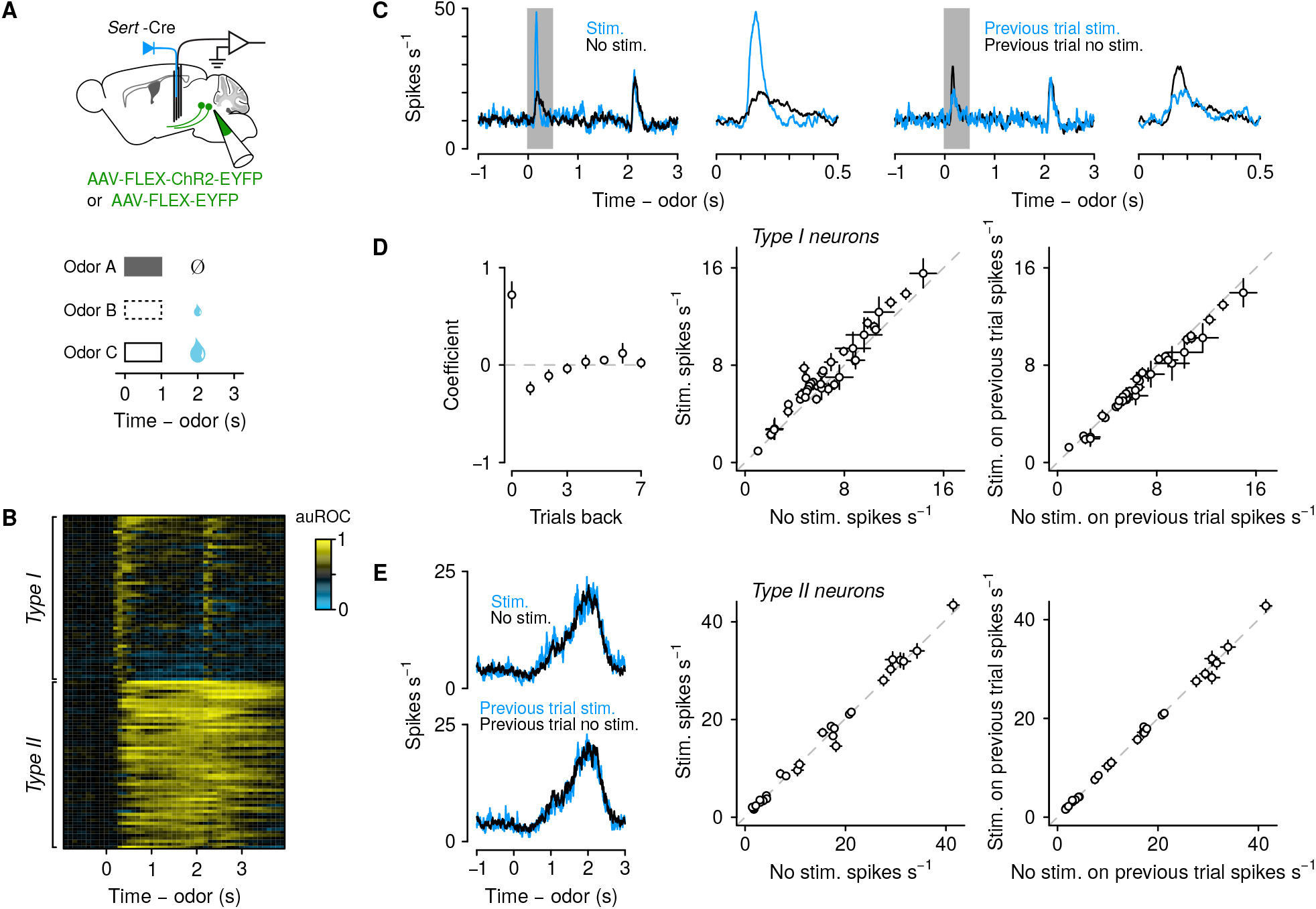
Stimulation of DR serotonergic axons in VTA mimics reward prediction errors in putative dopaminergic neurons. (A) Schematic of experiment to record extracellularly from VTA neurons while stimulating serotonergic axons in VTA (top) during a classical conditioning task (bottom). (B) Area under the receiver operating characteristic curve (auROC) during big-reward trials, for firing rates in 100-ms bins relative to baseline (pre-cue). Neurons are classified as *type I*, showing brief excitation at cues and rewards, or *type II*, showing sustained excitation following reward-predictive cues.

We recorded extracellularly from 111 VTA neurons in 8 *Slc6a4* -Cre (also known as *Sert* -Cre) mice, using multiple tetrodes (Figure 5A). In 4 of these mice, we injected adeno-associated virus (AAV) to express the light-gated ion channel, channelrhodopsin-2 (ChR2), specifically in serotonergic neurons (AAV-DIO-ChR2-EYFP). In the other 4 mice, as a control experiment, we expressed only a fluorescent protein (AAV-DIO-EYFP). During 30% of trials chosen at random, we stimulated serotonergic axons in VTA during the CS interval.

We first verified the presence of two types of neurons previously defined in VTA recordings in this task: *type I* neurons showed brief excitation at CS or reward, whereas *type II* neurons showed sustained excitation following reward-predictive CS (Cohen et al., 2012; Eshel et al., 2015). We replicated this observation here, by calculating the area under the receiver operating characteristic curve (auROC), comparing firing rates during large-reward trials to their baseline activity (pre-CS). auROC values close to 1 indicate increases in firing rate, values close to 0 indicate decreases in firing rate, and values of 0.5 indicate no modulation. We found 55 neurons classified as *type I* and 44 classified as *type II* (Figure 5B; a small number of neurons fell into the previously-defined *type III* cluster, but we do not consider them further here). In previous work, identified dopaminergic neurons were classified as *type I*, and identified GABAergic neurons were classified as *type II* (Cohen et al., 2012).

Of the 55 *type II* neurons, 39 were recorded in mice with ChR2-EYFP, and 16 in mice with EYFP alone. To quantify the effects of stimulating serotonergic axons on nearby VTA neurons, we calculated spike density functions, plotting the average firing rate of neurons on trials with or without stimulation. The example neuron shown in Figure 5C shows an increase in firing rate during the stimulation relative to unstimulated responses. We next examined the responses on trials following stimulation. The same neuron showed a decrease in firing rate when the previous trial included stimulation, relative to when the previous trial was unstimulated.

To quantify these effects across the population of 55 *type I* neurons, we calculated a regression, estimating the effect of serotonergic axonal stimulation history on the responses of VTA neurons to CS. In this model, the dependent variables were firing rates in the 1-s CS period, and regressors were the presence or absence of stimulation in the previous 8 trials (Figure 5D). We found two effects, one on short timescales (during the stimulation, occurring over hundreds of milliseconds), and one on long timescales (previous trials, occurring over tens of seconds). Regression coefficients corresponding to stimulation on the current trial were significantly larger than 0 (*t*_38_ = −5.4, *p <* 0.0001), while regression coefficients corresponding to stimulation on the previous trial were significantly smaller than 0 (*t*_38_ = 5.4, *p <* 0.001). Firing rates across the population were larger during serotonergic axon stimulation than in its absence (paired *t*_38_ = 5.4, *p <* 0.0001). On the trials following stimulation, firing rates during the CS were significantly smaller than following no stimulation (paired *t*_38_ = 3.6, *p <* 0.01). We note that the shape of this trial-history function matches qualitatively that of VTA dopamine neuron reward prediction errors (Bayer and Glimcher, 2005). Recordings from *type I* neurons in mice expressing EYFP alone in serotonergic neurons showed no significant modulation of firing rates during stimulation (*t*_15_ = −1.6, *p >* 0.55) or on subsequent trials (*t*_15_ = 5.4, *p <* 0.19).

These effects were cell-type specific. We calculated firing rates of *type II* neurons, with and without stimulation (Figure 5E). Of the 44 *type II* neurons, 28 were recorded in mice with ChR2-EYFP, and 16 in mice with EYFP alone. We counted spikes in the interval between CS onset and the end of the delay period (a total of 2 s). Regression coefficients corresponding to stimulation on the current or previous trials were not significantly different from 0 (for ChR2-EYFP experiments, current trial: *t*_27_ = 0.78, *p >* 0.4; previous trial: *t*_27_ = 0.097, *p >* 0.9). Likewise, spike counts during stimulation were not significantly different from those without on the current (paired *t*_27_ = 0.90, *p >* 0.3) or previous (*t*_27_ = 0.51, *p >* 0.6) trial (Figure 5E). Thus, the observed effects of serotonin neuron stimulation on putative dopamine neurons did not appear to require nearby putative GABA neurons.

## Discussion

Understanding heterogeneity among VTA dopamine neurons requires multiple lines of evidence: distributions of transcriptional expression, afferents and efferents, and physiological responses to behavioral variables. Here, we show that identified medial versus lateral VTA dopamine neurons are characterized by differential densities of molecular subtypes of neurons, and that they receive inputs from different populations of DR serotonin neurons. In addition, serotonin neuron axon stimulation affects putative dopamine neurons in lateral VTA on multiple timescales: excitation during the stimulation, and inhibition on trials following stimulation (several seconds later). The effects of stimulation may be independent of nearby VTA GABAergic neurons: we found no evidence of axonal stimulation on responses of functionally-defined *type II* neurons.

Our anatomical tracing data suggest an exquisite topographical organization of serotonergic projections to VTA, although the details of DR projections to other forebrain structures (Soubrié et al., 1984) and neurochemical phenotypes (Cragg et al., 1997; Fu et al., 2010) appear to vary across species. In mice, there appears to be a clear medial-lateral topography: medial DR serotonin neurons, many of which may co-release glutamate, target medial subdivisions of VTA, while lateral DR serotonin neurons (primarily non-glutamatergic) target lateral VTA. The medial-to-lateral gradient in DR projections corresponds to a gradient from mixed glutamatergic and serotonergic inputs to pure serotonergic inputs.

One possibility is that this is matched to a corresponding gradient in the expression of postsynaptic serotonin and glutamate receptors in dopamine neurons. We found evidence to support differential expression of glutamate receptors characterized by an enhancement of *Grm1/5* metabotropic receptors in a lateral dopaminergic subtype and enhanced *Grin2a* expression in a medial subtype. In general, these differences are consistent with a bias toward more rapid glutamate receptor kinetics (*Grin2a*-containing NMDA receptors are faster than *Grin2c*-containing; Cull-Candy and Leszkiewicz, 2004) in medial dopamine neurons. Serotonin receptor transcript levels were in general low in dopamine neurons with the highest average expression of *Htr2* isoforms; however, there were biases in the fraction of dopamine subtypes expressing *Htr3a* and *Htr7*. In contrast to glutamate receptors, these differences are more consistent with faster serotonergic effects in lateral VTA Sox6+ dopamine neurons as compared with medial VTA Tacr3+ subtypes. We have only examined a small fraction of the features of dopaminergic neuron diversity with this new single-cell sequencing dataset that is distinct in several ways (SMRT-Scrb sequencing from picked Slc6a3+ dopamine neurons) and will complement existing publicly available characterizations of the molecular diversity of dopamine neurons.

Previous literature showed multiple effects of serotonergic inputs onto dopaminergic neurons. Many studies focused on the immediate effects (i.e., during stimulation of DR). In *ex vivo* slice recordings, serotonin increases the inhibitory effect of dopamine onto dopaminergic neurons (Brodie and Bunney, 1996), and reduces I_h_ via 5-HT_2_receptors (Liu et al., 2003). Previous studies reported increases in dopamine concentrations in nucleus accumbens following serotonin infusion into VTA (Guan and McBride, 1989) or serotonergic axon stimulation in VTA (Wang et al., 2019). *In vivo* electrophysiological studies in anesthetized rats found that DR stimulation evoked a mixture of excitation (via 5-HT_1_A receptors; Prisco et al., 1994; Lejeune and Millan, 1998) and inhibition (Prisco et al., 1994; Gervais and Rouillard, 2000). However, other studies found inhibitory effects of serotonin on dopamine. Serotonin reuptake inhibitors decrease firing rates of putative dopaminergic neurons in anesthetized rats (Prisco and Esposito, 1995).

By recording from VTA neurons while stimulating their serotonergic afferents from DR, we revealed a straight-forward, cell-type-specific effect: short-term excitation, followed by delayed inhibition. The shape of this biphasic response mimics in form—and timescale—prediction error calculation in dopamine neurons. We propose that serotonergic inputs may contribute to reward prediction error calculations in dopamine neurons. These inputs could be used directly, as part of the normal calculation of reward prediction errors (Tian et al., 2016), or as a slower signal used to modulate the magnitude of reward prediction errors by changing learning rate or reward rate (Daw et al., 2002; Daw and Touretzky, 2002; Doya, 2002; Tobler et al., 2005; Doya, 2008; Matias et al., 2017; Grossman et al., 2020). Alternatively, the small increase in dopamine neuron firing rates during serotonergic stimulation could induce a negative reward prediction error, reflected in the next trial’s decreased CS response.

We found that effects of stimulating serotonergic inputs differed by VTA cell type. In particular, effects on *type I* neurons (which comprised dopaminergic neurons in previous studies) did not require concomitant effects on *type II* (which comprised GABAergic neurons in previous studies). We suggest that serotonin neuron modulation of dopamine neurons may involve direct synapses from one neuromodulator to the other. One limitation of this experiment is that we focused on lateral VTA recordings, given their well-studied response properties in single unit recordings. Medial VTA dopamine neurons are understudied; it will be important for future studies to determine their responses across behavioral tasks. Population calcium imaging from medial VTA has revealed important differences in response properties (de Jong et al., 2019; Cai et al., 2020); however, our data in lateral VTA suggest that electrophysiological measurements from single neurons (yet to be described in very medial VTA) may be critical to dissect the contributions of mixed serotonin and glutamate release, particularly given the heterogeneity of receptors expressed in VTA. Ultimately, understanding the function of serotonergic inputs to dopamine neurons will require using behavioral tasks in which the relevant variables vary over multiple timescales.

## Materials and methods

### Anatomical tracing experiments

#### Animals

All procedures were conducted in accordance with the National Institutes of Health Guide for the Care and Use of Laboratory Animals and approved by the Johns Hopkins University Animal Care and Use Committee. Eight C57BL/6J male mice (The Jackson Laboratory, 000664), aged 10–16 weeks at the time of first surgery (18–28 g), were housed with littermates in a reverse 12 hour-dark/12 hour-light cycle room (lights on at 20:00) with *ad libitum* food and water.

#### Surgeries

Mice were placed under isoflurane anesthesia (1.0–2.0% in O_2_ at 0.6–1.0 L/min) for stereotactic surgery under aseptic conditions. 40–70 nl retrobeads (LumaFluor; red beads) were injected with a glass pipette and manual injection pump (MMO-220A, Narishige) at a rate of 20 nl/min. Injections to target the VTA were aimed at the following coordinates relative to bregma: anterior-posterior −3.3 mm; dorsal-ventral +4.5 mm; medial-lateral +0.3 mm (most medial injections) through +0.7 mm (most lateral injections). Glass pipettes were left in place for 15 min following full injection volume delivery. Five to 7 days after retrobead injections (to allow for sufficient retrograde transport), animals were intracerebroventricularly injected with 300 nl colchicine dissolved in physiological saline (6 mg/ml) per ventricle at a rate of 75 nl/min. Injections to target bilateral ventricles were aimed at the following coordinates relative to bregma: anterior-posterior +0.3 mm; medial-lateral *±*1.0 mm; dorsal-ventral +3.0 mm. Colchicine injections allowed for better visualization of the vesicular protein VGLUT3 in somata. All surgeries were followed by analgesia (ketoprofen: 5 mg/kg, subcutaneous; buprenorphine: 0.1 mg/kg, subcutaneous) and postoperative care (ketoprofen administration and 500 ml subcutaneous physiological saline injection) for 2 days.

#### Immunohistochemistry

One day following colchicine injection, mice were deeply anesthetized (3% in O_2_ at 1.0 L/min) and perfused with 4% paraformaldehyde dissolved in phosphate buffer. Brains were post-fixed for 16 hr, switched to phosphate buffer, and sectioned one to two weeks later. Coronal sections were prepared at 50 *µ*m in phosphate buffer solution (PBS) on a vibratome. Free floating sections were washed three times with PBS and moderate shaking for 10 min per wash. Sections were then permeabilized for 1 hr (0.5% TritonX in PBS) at room temperature, followed by blocking for 1 hr (0.5% TritonX in 10% Fetal Bovine Serum in PBS) at room temperature. Sections were incubated with primary antibodies diluted in blocking solution (see region-specific details below) overnight at 4°C with mild shaking. Following overnight incubation, sections were washed once with wash buffer (0.05% Tween-20 in PBS) for 15 min, then twice with PBS for 10 min per wash. Sections were then incubated with secondary antibodies diluted in blocking solution (see region-specific details below) overnight at 4°C with mild shaking. Following overnight incubation, sections were again washed once with wash buffer for 15 min, then twice with PBS for 10 min per wash. Tissue was then mounted on glass slides with DAPI-mounting media (Vector Laboratories: Vectashield Antifade Hard Set Mounting Medium with DAPI, to visualize nuclei) and imaged within two weeks. Region-specific information: DR sections (from −5.0 mm to −4.0 mm from bregma in the anterior-posterior axis) were co-stained with primary antibodies against tryptophan hydroxylase-2 (goat *α*-TPH2, Abcam 121020, at 1:400) as a marker of serotonergic neurons and vesicular glutamate transporter-3 (guinea pig *α*-VGLUT3, Synaptic Systems 135-204, at 1:400) as a marker of glutamatergic neurons that express VGLUT3. We tested the VGLUT3 antibody in a Vglut3 knockout mouse, Slc17a8^tm1Edw^ (Seal et al., 2008), and confirmed its specificity. Secondary antibodies used to visualize the protein expression were donkey *α*-goat conjugated to Cy5 (Abcam ab6566) and donkey *α*-guinea pig conjugated to Alexa Fluor 488 (Jackson ImmunoResearch 706-545-148).

#### Image acquisition

VTA sections were first imaged at 2*×* to confirm and localize retrobead injection sites in reference to ventral midbrain dopamine neurons. Sections of the DR were imaged (Zeiss Confocal Laser Scan Microscope 800 with Airyscan) at 20*×*, scan speed 7, resolution 512 *×* 512 per tile, with Z-stack and tiling (4 *×* 4) for visualization of DAPI (405 nm), VGLUT3 (488 nm), retrobeads (594), and TPH2 (640 nm).

#### Data analysis

VTA injection sites were reconstructed by standardizing the −3.3 mm section across all animals. Reconstruction points marked the bottom (dorsal-ventral axis) and center (medial-lateral axis) of injection. Actual injection reconstructions were created by taking fluorescent images with the same settings and then vectorizing the image with the same brightness threshold to maintain consistent comparisons across mice. DR neurons were manually counted using Fiji (Schindelin et al., 2012) with the Bio-Formats and Cell Counter plugins. Neurons were identified as being one type in four groups: 1) TPH2+/VGLUT3− 2) TPH2-/VGLUT3+ 3) TPH2+/VGLUT3+ or 4) TPH2−/VGLUT3−. Coordinates of all marked neurons were extracted using a custom written script in Python (Version 3.6) and standardized to a manually-selected reference point (ventromedial point of the cerebral aqueduct) to systematically compare across mice and the anterior-posterior axis. Statistical analyses were conducted using Python packages Matplotlib, Seaborn, SciPy, or scikit-learn.

### Electrophysiology and behavior

#### Animals

All procedures were conducted in accordance with the National Institutes of Health Guide for the Care and Use of Laboratory Animals and approved by the Johns Hopkins University Animal Care and Use Committee. Eight heterozygous *Slc6a4* -Cre (also known as *Sert* -Cre; Zhuang et al., 2005) male mice (18–28 g), aged 8–16 weeks at time of first surgery, were housed with littermates in a reverse 12 hr-dark/12 hr-light cycle room (lights on at 8:00 pm) with *ad libitium* water and food.

#### Surgeries

Four mice (experimental group) were stereotactically injected in the DR with AAV2/5-EF1*α*-DIO-ChR2-WPRE-eYFP (Addgene viral prep 20298-AAV5; a gift from Karl Deisseroth) and four mice (control group) were stereotactically injected in the DR with AAV2/5-EF1*α*-DIO-WPRE-eYFP (Addgene viral prep 27056-AAV5) at a rate of 75 nl per min using surgical procedures (anesthesia, analgesia, and postoperative care) described above. Injections to target the dorsal raphe were aimed at the following five coordinates relative to bregma: 1) anterior-posterior −4.6 mm; medial-lateral −0.2*mm*; dorsal-ventral +2.9 mm; 2) anterior-posterior -4.6 mm; medial-lateral +0.2 mm; dorsal-ventral +2.9 mm; 3) anterior-posterior −4.6 mm; medial-lateral −0.2 mm; dorsal-ventral +3.3 mm; 4) anterior-posterior −4.6 mm; medial-lateral +0.2 mm; dorsal-ventral +3.3 mm; 5) anterior-posterior −4.6 mm; medial-lateral 0.0 mm; dorsal-ventral +3.1 mm, with a volume of 100 nl per injection site. Following two weeks from the virus injections, a custom-built headplate was affixed to the skull (C&B-Metabond, Parkell) and custom-built advanceable (dorsal-ventral axis) microdrives were unilaterally implanted under surgical procedures (anesthesia, analgesia, and postoperative care) described above into the VTA (relative to bregma: anterior-posterior −3.1 mm – −3.3 mm; medial-lateral +0.6 mm; dorsal-ventral +4.0 mm). Microdrives were built with eight nichrome wire tetrodes (PX000004, Sandvik) positioned inside 39 ga polyimide guide tubes and glued around a 200 *µ*m optic fiber (0.39 NA). Wires were cut to be 300–500 *µ*m below the bottom of the fiber and were gold-plated to impedances between 250–350 kΩ.

#### Behavior

Four weeks after surgery (to allow for animal recovery and sufficient protein expression in axon terminals), mice were water restricted to 80% of their body and began behavioral training. Mice were first habituated to the behavioral rig and were allowed free access around the rig for 20 min for the first day. For the second, third, and fourth day, mice were head-fixed and given 1 ml of water (through the reward tube) spread across 10 min, 20 min, and 30 min, respectively. On the fifth day, mice were put through the classical conditioning behavioral task during the dark phase and at the same time of the day (approximately 1 hr session between 09:00 and 15:00). Each behavioral session consisted of 400 trials, in which three different odors (limonene, citral, and pentyl acetate diluted in mineral oil at 1:10, counterbalanced across animals) were pseudo-randomly chosen and delivered for one second with a custom-built olfactometer (Bari et al., 2019). Each odor was associated with an outcome two seconds following the onset of odor delivery: no reward, small reward (1.5 *µ*l of water delivered through a 27 ga stainless steel tube), or large reward (4.0 *µ*l of water). A variable inter-trial interval (at least 2 s), was chosen from an exponential distribution with a mean of approximately 6 s. Following 11 days of behavioral task training (not including the four habituation days), the laser stimulation protocol began in which light (473 nm, 50 Hz, approximately 5 mW, 1 s) was delivered to optogenetically activate serotonin axons concurrent with the odor presentation in 30% of pseudo-randomly selected trials (across all three trial types). Mice typically drank 500–700 *µ*l of water in a daily session and were supplemented to receive a total of 1 ml per day.

#### Electrophysiology

Extracellular signals of neurons dorsal to or in VTA were recorded while the animals performed the behavioral task. After each behavioral session, microdrives were ventrally advanced by 30 m, which allowed for recordings to be in ventral tegmental area once the laser stimulation protocol began. Extracellular signals were recorded at 32 kHz and broadband filtered between 0.1 and 9,000 Hz (Neuralynx). Single units were spike-sorted using two dimensional projections of spike amplitude. *Type I* neurons were classified using the following criteria (Cohen et al., 2012): 1) monotonic encoding of forthcoming reward size during the odor presentation period; and 2) low baseline firing rate (*<* 15 spikes per second). *Type II* neurons were classified using the following criteria: 1) recorded in the same session as *type I* neurons; and 2) persistent, sustained excitation during the delay between odor and reward that scaled monotonically with forthcoming reward size. Spike sorting and further analyses were performed in R, Python, and Matlab. Statistical tests are reported with Bonferroni corrections for multiple comparisons.

### Single-cell sequencing

#### Animals

All animal procedures were performed in accordance with protocols approved by the Institutional Animal Care and Use Committee at the Janelia Research Campus and consistent with the standards set forth by the Association for Assessment and Accreditation of Laboratory Animal Care. For single-cell transcriptomics, 2 male (24.0 g – 24.1 g) and 2 female (18.6 g – 18.8 g) *Slc6a3* -Cre*×*Ai14 (Bäckman et al., 2006; Madisen et al., 2010) littermate mice were used. For representative histology, 1 male (24.0 g) Scl6a3-Cre::Ai14 was used. All mice were 8–9 weeks old at the time of the first experiment. Mice were group-housed on a 12-hr reverse light/dark cycle (08:00 – 20:00 dark) with food and water available *ad libitum*.

#### Surgeries

For labeling of the lateral VTA, 100 *µ*l retrobeads were injected at an injection rate of 100 *µ*l/min bilaterally into the lateral shell of the nucleus accumbens (NAc; Lammel et al., 2008) using glass pipettes. Coordinates relative to bregma were as follows: anterior-posterior: +0.98 mm; medial-lateral: 1.80 mm; dorsal-ventral: −4.92 mm. All surgeries were performed stereotactically and aseptically, with mice anesthetized (1.0 – 2.0% isoflurane in O_2_ at 0.6 – 1.0 L/min). Prior to surgery, mice were administered with bupinorphine (0.1 mg/kg). Following surgery, mice were placed on a heating pad (37°C) for two hours and were administered with ketoprofen (5 mg/kg) for three days. The time between retrobead injection to cell collection ranged from 6–29 days.

#### Single-cell collection and sequencing

For tissue processing and single-cell collection, mice were deeply anesthetized (3.0% isoflurane in O_2_ at 0.6–1.0 L/min) and rapidly decapitated. Coronal slices were sectioned at 300 *µ*m on a vibratome (Leica VT1200S, Leica Microsystems) and incubated for 1 hour at room temperature in artificial cerebrospinal fluid (ACSF) containing TTX. Slices around the NAc (+2 mm through −1 mm from bregma in the anterior-posterior axis) were first visualized under the fluorescent dissecting microscope at 1*×* – 5*×* to confirm accurate bilateral injections into the lateral shell of the NAc. If one side was not accurately targeted, only the ipsilateral side was used for cell collection. If both sides were not accurately targeted, the mouse was not used for cell collection. Slices containing ventral midbrain dopamine neurons were confirmed with visualization of tdTomato+ neurons. Generally, two slices were used: −3.0 mm through −3.3 mm and −3.3 mm through −3.6 mm from bregma in the anterior-posterior axis. Cuts to separate medial VTA and the lateral VTA were made under the dissecting microscope. Medial VTA extended from the midline to the medial boundary of the green retrobeads. Lateral VTA was determined by the boundaries of the lateral shell NAc-projecting neurons as well as the axon bundle separating VTA and the substantia nigra. Following microdissection, the tissue was dissociated, washed 3 times with ACSF, and transferred to clean dishes. Individual dopamine cells were visualized under the fluorescent dissecting microscope, picked out with aspiration, placed in single test tubes containing lysis buffer here, and stored at −80°C until sequencing.

#### Sorted single-cell RNA-seq

Samples for single-cell RNA-seq were prepared as previously described (Cembrowski et al., 2018). Briefly, acute brain slices were cut from brains of *Slc6a3* -Cre *×* Ai9 mice (*n* = 4 male; *n* = 4 female) that had been injected with green fluorescent retrograde beads (Lumiflor) injected into the NAc. The VTA was identified in coronal sections containing red fluorescent dopamine neurons and sections containing retrograde beads were selected. Three subregions within the midbrain were made by two cuts prior to tissue dissociation and single cell isolation. One cut was made to separate substantia nigra from lateral VTA and a second cut was made to isolate medial VTA from lateral VTA. This cut was chosen such that lateral VTA tissue contained all retrogradely labelled cells. Cut sites are schematized on an example section in Figure 1A. Brain sections including the VTA were digested with 1 mg/ml protease (Sigma-Aldrich). Single cells from each of these three subregions were then obtained from dissociated tissue from medial and lateral VTA and collected separately into 8-well strips containing 3 *µ*l Smart-Seq2 lysis buffer, flash-frozen on dry ice and stored at −80^*°*^C until further use (Picelli et al., 2013).

After thawing, cells were lysed and digested with Proteinase K, and 1 *µ*l of ERCC RNA spike-in mix at 10:7 dilution (Life Technologies) and barcoded reverse transcription (RT) primers were added. cDNA was synthesized using Maxima H Minus Reverse Transcriptase (Thermo Fisher Scientific) and E5V6NEXT template switch oligonucleotide, followed by heat inactivation of reverse transcriptase. PCR amplification using a HiFi PCR kit (Kapa Biosystems) and SINGV6 primer was performed with a modified thermocycling protocol (−98^*°*^C for 3 min, 20 cycles of −98^*°*^C for 20 s, −64^*°*^C for 15 s, −72^*°*^C for 4 min, final extension at −72^*°*^C for 5 min). Samples were then pooled across strips, purified using Ampure XP beads (Beckman Coulter), washed twice with 70% ethanol and eluted in water. These pooled strips were then combined to create the plate-level cDNA pool for tagmentation, and concentration was determined using the Qubit High-Sensitivity DNA kit (Thermo Fisher Scientific).

Tagmentation and library preparation using 600 pg of cDNA from each plate of cells was then performed with a modified Nextera XT (Illumina) protocol, but using the P5NEXTPT5 primer and tagmentation time extended to 15 min (Soumillon et al., 2014). The libraries were then purified following the Nextera XT protocol (at 0.6*×* ratio) and quantified by qPCR using Kapa Library Quantification (Kapa Biosystems). A total of 6–10 plates were run on a NextSeq 550 high-output flow cell. Read 1 contained the cell barcode and unique molecular identifier (UMI). Read 2 contained a cDNA fragment from the 3’ end of the transcript.

### Single-cell RNA-seq analysis

#### Data processing and quality control

Single-cell RNA-seq data were trimmed for adapters using cutadapt and aligned to the mouse genome (mm10) using STARsolo (https://github.com/alexdobin/STAR/blob/master/docs/STARsolo.md) based upon the STAR aligner (Dobin et al., 2013). After preprocessing, single cells were required to have more than 30,000 UMIs and more than 2,000 genes detected per cell, which yielded a total of 722 single cells. 361 cells were obtained from medial VTA tissue and 342 from lateral VTA tissue. Of these, 16 cells were found to have significantly low counts of Slc6a3 (suggesting possible contamination given collection from *Slc6a3* -Cre::Ai9 mice), however, inclusion did not substantially affect any results and thus our analysis contains these cells for completeness. Previous evaluation of this same method through quantification of ERCC spike-in control RNA indicated high accuracy and sensitivity of our single-cell profiling (Phillips et al., 2019).

#### Histology and imaging

Slices (300 *µ*m) containing retrobead injection sites in the lateral shell of the NAc were mounted (Vector Laboratories VECTASHIELD Hardset Antifade Mounting Medium) and cover-slipped following injection confirmation under the fluorescent microdissection microscope. Images were acquired with a 4*×* objective.

## Acknowledgements

We thank T. Shelley for machining. J.T.D. is a Senior Group Leader at the Janelia Research Campus (JRC) of the Howard Hughes Medical Institute. This work was supported by Klingenstein-Simons, MQ, NARSAD, Whitehall, R01DA042038, and R01NS104834 (J.Y.C.), and P30NS050274. Part of this work was supported by the Visiting Scientist Program at JRC.

## Author contributions

A.J.C., L.W., and A.L. performed single-cell sequencing. A.J.C. and J.T.D. analyzed sequencing data. A.J.C., F.L., and M.A. collected behavioral, electrophysiological, and retrograde tracing data. A.J.C. and J.Y.C. analyzed behavioral, electrophysiological, and retrograde tracing data. A.J.C., J.T.D., and J.Y.C. designed the study and wrote the paper.

---

**Figure S1.**
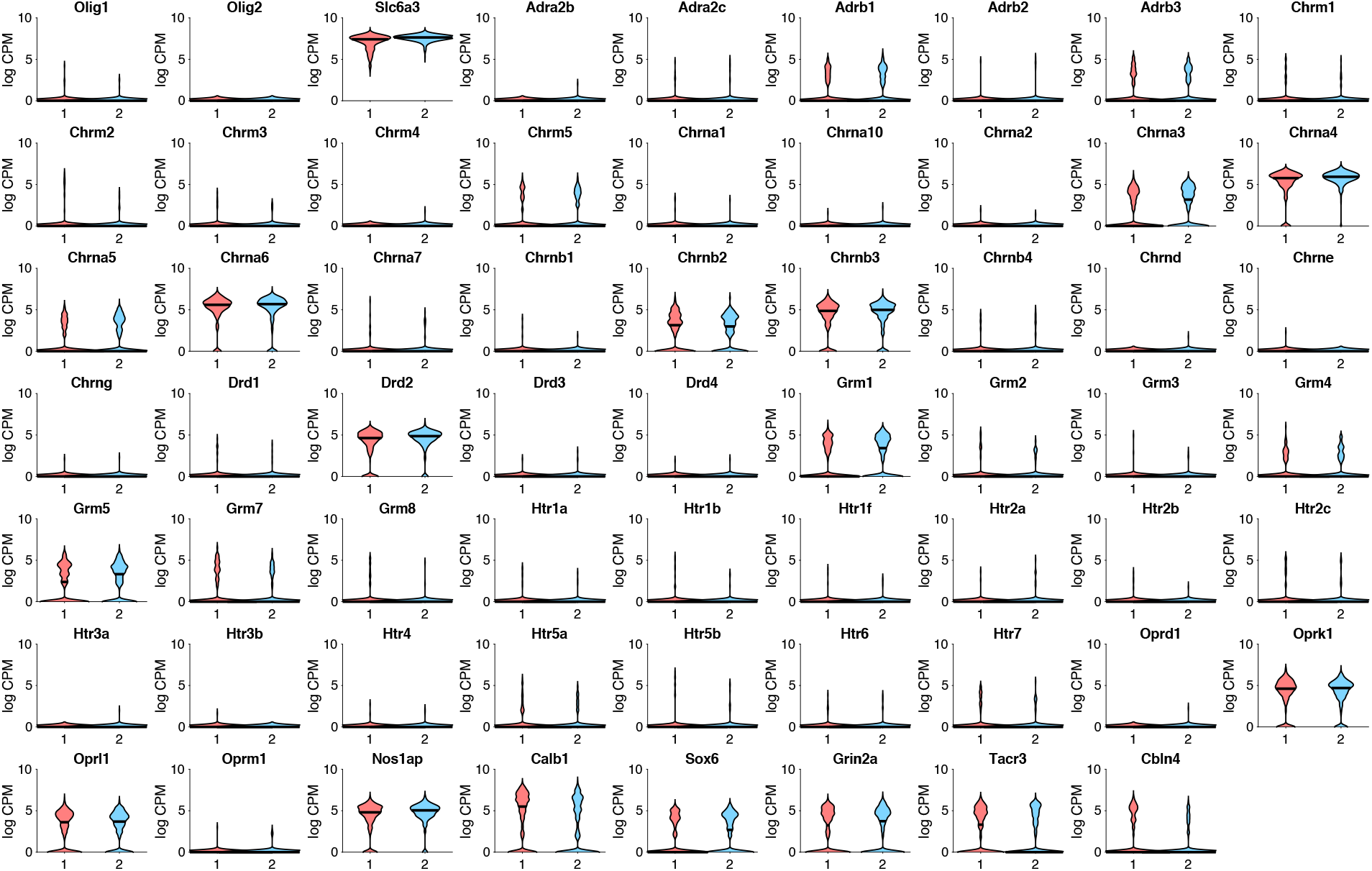
Additional transcripts from medial and lateral VTA dopamine neurons.

